# AMELIE 2 speeds up Mendelian diagnosis by matching patient phenotype & genotype to primary literature

**DOI:** 10.1101/839878

**Authors:** Johannes Birgmeier, Maximilian Haeussler, Cole A. Deisseroth, Ethan H. Steinberg, Karthik A. Jagadeesh, Alexander J. Ratner, Harendra Guturu, Aaron M. Wenger, Mark E. Diekhans, Peter D. Stenson, David N. Cooper, Christopher Ré, The Manton Center, Alan H. Beggs, Jonathan A. Bernstein, Gill Bejerano

## Abstract

The diagnosis of Mendelian disorders requires labor-intensive literature research. Trained clinicians can spend hours looking for the right publication/s supporting a single gene that best explains a patient’s disease. AMELIE (<u>A</u>utomatic <u>Me</u>ndelian <u>Li</u>terature <u>E</u>valuation) greatly accelerates this process. AMELIE parses all 29 million PubMed abstracts, downloads and further parses hundreds of thousands of full text articles in search of information supporting the causality and associated phenotypes of any published genetic variant. AMELIE then prioritizes patient candidate variants for their likelihood of explaining any patient’s given set of phenotypes. Diagnosis of singleton patients (without relatives’ exomes) is the most time-consuming scenario. AMELIE ranked the causative gene at the very top in 2/3 of 215 diagnosed singleton Mendelian patients. Evaluating only the top 11 AMELIE scored genes of 127 (median) candidate genes per patient results in rapid diagnosis for 90+% of cases. AMELIE-based evaluation of all cases is 3-19x more efficient than hand-curated database-based approaches. We replicate these results on a cohort of clinical cases from Stanford Children’s Health and the Manton Center for Orphan Disease Research. An analysis web portal with our most recent update, programmatic interface and code will be available at AMELIE.stanford.edu. A pilot run of the web portal has already served many thousands of job submissions from dozens of countries.

## Introduction

Millions of babies are born worldwide each year affected by severe genetic and often Mendelian disorders *(1)*. Patients with Mendelian diseases have one or two genetic variants in a single gene primarily responsible for their disease phenotypes *(2)*. Roughly 5,000 Mendelian diseases, each with a characteristic set of phenotypes, have been mapped to about 3,500 genes to date *(3)*. To identify candidate causative genes, exome sequencing is often performed, with a relatively high (currently 30%) diagnostic yield *(4)*. A genetic diagnosis provides a sense of closure to the patient family, aids in patient trajectory prediction and management, allows for better family counseling, and in the age of gene editing even provides first hope for a cure. However, identifying the causative mutation/s in a patient’s exome to arrive at a diagnosis can be very time-consuming, with a typical exome requiring hours of expert analysis *(5)*.

Definitive diagnosis of a known Mendelian disorder is accomplished by matching the patient’s genotype and phenotype to previously described cases from the literature. Manually curated databases *(6–10)* are utilized to more efficiently access extracts of the unstructured knowledge in the primary literature. Automatic gene ranking tools *(11–18)* use these databases to prioritize candidate genes in patients’ genomes for their ability to explain patient phenotypes. An important feature of many gene ranking tools is the use of phenotype match functions on patient phenotypes and gene/disease-associated phenotypes. Phenotype match functions exploit the structure of a phenotype ontology *(9)* and known gene-disease-phenotype associations to quantify the inexact match between two sets of phenotypes *(11, 12)*, with recent approaches developed to computationally extract phenotype data from electronic medical notes *(19, 20)*. The goal of all gene ranking tools is to aid a busy clinician in arriving at a definitive diagnosis of any case presented to them in the shortest amount of time, by reading up on genes in the order the algorithm has ranked them.

Given the rapidly growing number of rare diseases with a known molecular basis *(21)* and the difficulty of manually finding a diagnosis for some rare diseases with variable phenotypes, many patients experience long diagnostic odysseys *(22)*. Expert clinician time is expensive and scarce, but machine time is cheap and plentiful. We aim to accelerate the diagnosis of patients with Mendelian diseases by using information from primary literature to construct gene rankings, thus allowing clinicians to discover the causative gene along with supporting literature in a minimum amount of time.

Here, we introduce AMELIE (<u>A</u>utomatic <u>Me</u>ndelian <u>Li</u>terature <u>E</u>valuation). AMELIE uses natural language processing (NLP) to automatically construct a homogeneous knowledgebase about Mendelian diseases directly from primary literature. To perform this operation, AMELIE is trained on data from manually curated databases such as OMIM *(6)*, HGMD *(8)* and ClinVar *(10)*. AMELIE then uses a machine learning classifier that integrates knowledge about a patient’s phenotype and genotype with its knowledgebase to rank candidate genes in the patient’s genome for their likelihood of being causative, and simultaneously supports its ranking results with annotated citations to the primary literature. We show that this end-to-end machine learning approach outperforms gene ranking methods that rely on manually curated databases using a total of 271 singleton patients from 3 different sources (including 2 clinical centers and a research cohort).

## Methods

### Overview of AMELIE

Given a patient’s genome sequencing data and a phenotypic description of the patient, AMELIE aims to both identify the gene causing the patient’s disease (when possible) and supply the clinician with literature supporting the gene’s causal role. To this end, AMELIE creates a ranking of candidate causative genes in the patient’s genome with the aim of ranking the true causative gene at the top. AMELIE constructs its candidate causative gene ranking by comparing information from the primary literature to information about the patient’s genotype and phenotype.

In order to process information from the full text of primary literature, AMELIE constructs a knowledgebase (the “AMELIE knowledgebase”) directly from the primary literature up-front using natural language processing techniques trained on manually curated databases. After knowledgebase construction, AMELIE ranks any patient’s candidate causative genes using a classifier (the “AMELIE classifier”), which compares knowledge from the AMELIE knowledgebase with phenotypic and genotypic information about the patient. AMELIE explains each gene’s ranking to the clinician by citing articles about this gene in the AMELIE knowledgebase.

### AMELIE parses all of PubMed to construct an integrated knowledgebase about Mendelian diseases

#### Identification and download of relevant articles based on all of PubMed

The first step towards building the AMELIE knowledgebase is discovering relevant primary literature. Of 29 million peer-reviewed articles deposited in PubMed, only a fraction is relevant for Mendelian disease diagnosis. We constructed a machine learning classifier that, given titles and abstracts of articles from PubMed, identifies potentially relevant articles for the AMELIE knowledgebase.

Machine learning classifiers take as input a numerical vector describing the input, called the “feature vector”. Here, we used a so-called TF-IDF transformation to convert input text into a feature vector (Supplementary Methods). The title/abstract document classifier is implemented as a Logistic Regression classifier, using the Python 3.7.0 machine learning library scikit-learn 0.20.0. Logistic regression transforms its output using the logistic sigmoid function to return a probability value which is then mapped into binary (positive/negative) decision making *(23)*.

Machine learning classifiers learn to classify an input as positive (relevant) or negative (irrelevant) by being exposed to a large number of labeled positive and negative examples (called the training set). OMIM *(6)* is an online database of Mendelian diseases, genes and associated phenotypes. HGMD *(8)* is a database of disease-causing mutations in the human genome. The training set for the title/abstract relevance classifier consisted of titles and abstracts of 56,479 Mendelian-disease-related articles cited in OMIM and HGMD as positive training examples, and 67,774 random titles and abstracts of PubMed articles (which are largely unrelated to Mendelian disease) as negative training examples.

Precision and recall are two standard measures of evaluating classifier performance. Precision measures the fraction of all inputs classified positive that are truly positive (here: relevant). Recall measures the fraction of truly positive inputs that are classified positive. Five-fold cross-validation (splitting all available labeled training data to include 80% in a training set and evaluating on the remaining 20%, 5 times in round-robin fashion) returned an average precision of 98% and an average recall of 96%.

All 28,925,544 titles and abstracts available in PubMed on September 30, 2018, were downloaded and processed by the document classifier. The classifier identified 578,944 articles as possibly relevant based on their PubMed title and abstract, of which we downloaded 515,659 (89%) full text articles directly from dozens of different publishers using the software PubMunch3 version 1.0.3 *(24)* (Supplementary Methods).

#### Building a structured database of information about Mendelian diseases from full text

From the full text of an article, multiple types of information are extracted. Gene mentions in full text were identified using lists of gene and protein names and synonyms from the HUGO Gene Nomenclature Committee (HGNC) *(25)*, UniProt *(26)* and the automatically curated PubTator *(27)* (an NCBI service combining gene mentions discovered by multiple previously published automatic gene recognition methods). AMELIE recognizes approximately 93% of disease-causing gene names (Supplementary Methods). However, through a combination of unfortunate gene synonyms (such as “FOR”, “TYPE”, “ANOVA”, or “CO2”), as well as genes mentioned only in titles of cited references, or interaction partners of causative genes, a median of 12 distinct gene candidates are discovered in each article (see Supplementary Table S1).

To discover the correct gene(s) around which the paper revolves, each distinct gene candidate extracted from an article receives a “relevant gene score” between 0 and 1 that gives a likelihood of the gene being important in the context of the article. Training data for the relevant gene classifier was obtained from OMIM and HGMD (Supplementary Methods). A total of 304,471 downloaded full-text articles contained at least one gene with a relevance score of 0.1 or higher. These articles, along with their above threshold scoring genes, form the AMELIE knowledgebase. Articles in the AMELIE knowledgebase contained a median of 1 gene with a relevant gene score between 0.1 and 1 (see Supplementary Table S1).

Further, genetic variants (e.g., “p.Met88Ile” or “c.251A>G”) were identified in the full text of each article and converted to genomic coordinates (chromosome, position, reference and alternative allele) using the AVADA variant extraction method *(28)*. A median of 3 distinct genetic variants were extracted from 123,073 full-text articles in the AMELIE knowledgebase.

Phenotype mentions were recognized in articles’ full text using a list of phenotype names compiled from Human Phenotype Ontology (HPO) *(9)*. By linking all genes with a relevant gene score of at least 0.5 in an article with all phenotypes mentioned in the same article, we arrived at a total of 872,080 gene-phenotype relationships covering 11,537 genes (Supplementary Figure S1).

Five scores between 0 and 1 were assigned to the full text of each article. Training data for all of these classifiers were obtained from OMIM and HGMD. A “full-text document relevance” score assesses the likely relevance of the article for the diagnosis of Mendelian diseases. A “protein-truncating” and a “non-truncating” score each give an assessment of whether the article is about a disease caused by protein-truncating (splice-site, frameshift, stopgain) or non-truncating (other) variants. A “dominant” and a “recessive” score each give an assessment of the discussed inheritance mode(s) in the article.

Precision and recall of full-text article information (relevant genes, extracted phenotypes, full-text article scores) varied between 74% and 96% (Supplementary Methods). All the data described in this section are entered into the AMELIE knowledgebase, keyed on the article they were extracted from (Figure 1a). The top journals from which the most gene-phenotype relationships were extracted are shown in Figure 1b and Supplementary Table S2.

**Figure 1.**
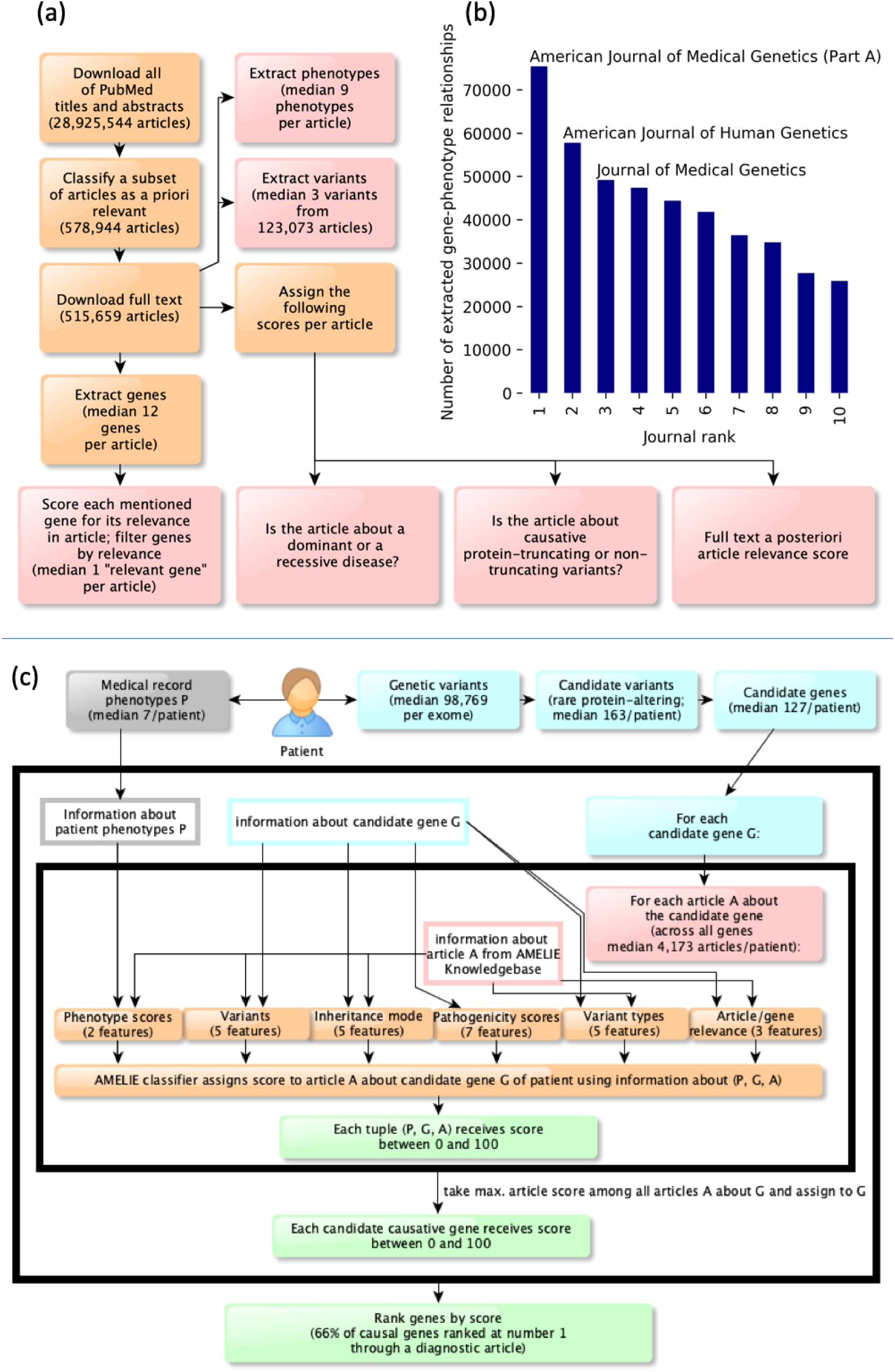
AMELIE knowledgebase creation and subsequent patient causal gene ranking classifier. **(a)** AMELIE knowledgebase creation. AMELIE applies multiple machine learning classifiers to all 29 million PubMed abstracts to parse, predict relevance, download full text, and finally extract Mendelian gene-phenotype relationships and related attributes automatically. **(b)** Number of gene-phenotype relationships extracted from top 10 journals. **(c)** The AMELIE classifier combines 27 features to rank all articles in the AMELIE knowledgebase for their ability to explain any input patient.

### AMELIE classifier assigns patient genes a likelihood of being causative

Given a patient with a suspected Mendelian disease, AMELIE aims to speed up discovery of the causative gene by ranking patient genes for their ability to describe a patient set of phenotypes.

#### Definition of candidate causative variants and genes

AMELIE performs standard filtering of the patient variant list *(21, 29)* to keep only “candidate causative variants” that are rare in the unaffected population and are predicted to change a protein-coding region (missense, frameshift, nonframeshift indel, core splice-site, stoploss, and stopgain variants). The core splice site is defined to consist of the 2 basepairs at either end of each intron. Genes containing candidate causative variants are candidate causative genes (or “candidate genes” for short). AMELIE can rank approximately 97% of known disease-causing mutations, excluding only those in deeper intronic and non-protein-coding intergenic regions (Supplementary Methods).

#### Per patient AMELIE gene-article scores

We define an article in the AMELIE knowledgebase to be *about* a candidate causative gene if the candidate causative gene has a relevant gene score of at least 0.1 in the article. We constructed a machine learning classifier called the “AMELIE classifier” that assigns a score between 0 and 100 to triples *(P, G, A)* consisting of a set of patient phenotypes *P*, a candidate causative gene *G*, and an article *A* about the candidate gene. Intuitively, given a patient with phenotypes *P* and a candidate gene *G*, the AMELIE score indicates whether the article *A* is likely helpful for diagnosing the patient because it links mutations in *G* to the patient’s phenotypes *P*. Higher AMELIE scores indicate more likely diagnostic articles. The AMELIE classifier is implemented as a scikit-learn version 0.20.0 *(30)* LogisticRegression classifier and returns a score between 0 and 100 called the “AMELIE score”. The AMELIE score is used to both rank patient candidate genes and explain rankings by citing primary literature, as described below.

#### Integration of information in the AMELIE knowledgebase with patient information to construct a feature set for the AMELIE classifier

The AMELIE classifier uses a set of 27 real-valued features, falling into 6 feature groups (Figure 1c). The 6 feature groups comprise: **(1)** 5 features containing information about disease inheritance mode extracted from the article and patient variant zygosity. **(2)** 5 features containing information about AVADA-extracted variants from the article and overlap of these variants with patient variants. **(3)** 2 features containing information about patient phenotypes based on the Phrank *(11)* phenotypic match score of phenotypes in the article *A* with the patient phenotypes *P*. **(4)** 5 features containing information about disease-causing variant types in the article and patient variant types. **(5)** 3 features containing information about article relevance and relevance of the candidate gene in the article, and **(6)** 7 features containing a-priori information about the patient’s candidate causative variants in *G* such as in-silico pathogenicity scores *(31)* and gene-level mutation intolerance scores *(32, 33)* (Supplementary Methods).

#### Training the AMELIE classifier to prioritize patient candidate causative genes

To train the AMELIE classifier, we constructed a set of 681 simulated patients using data from OMIM *(6)*, ClinVar *(10)* and the 1000 Genomes Project *(34)*. Each simulated patient *s* was assigned a disease from OMIM, with phenotypes noisily sampled from the phenotypes associated with the disease (Supplementary Methods). The genome of each simulated patient was based on genome sequencing data from the 1000 Genomes Project. An appropriate disease-causing variant from ClinVar was added to each simulated patient’s genome. Each simulated patient was assigned a diagnostic article *A*_*s*_ describing the genetic cause of the patient’s disease (Supplementary Methods). In total, the simulated patients cover a total of 681 OMIM diseases (1 per patient) and a total of 1,090 distinct phenotypic abnormalities (Supplementary Table S3). The sampled phenotypes for each disease cover an average 21% of the phenotypes that were manually associated with the disease by HPO.

The AMELIE classifier was trained to recognize the diagnostic article *A*_*s*_ out of all articles about genes with candidate causative variants in a patient *s*. Of a total of 681 training patients that were constructed using data in OMIM and ClinVar, the single positively labeled article was recognized and downloaded during AMELIE knowledgebase construction in 664 cases (98%), thus creating 664 positive training examples. The negative training set for the AMELIE classifier consisted of triples *(P*_*s*_, *G, A)* for each simulated patient *s* where *G* was a non-causative candidate gene in patient *s*, and *A* was an article about *G*. For training efficiency, we used only 664,000 random negative training examples out of all available negative training examples.

#### Ranking candidate causative genes using the AMELIE classifier

The AMELIE classifier assigns each candidate gene *G* the score of its highest-scoring triple *(P, G, A)* (Figure 1c):

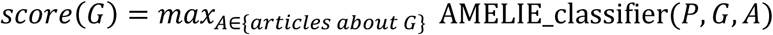

Candidate causative genes are ranked in descending order of their associated score.

## Results

### Evaluating AMELIE on a real patient test set

We evaluated AMELIE on a set of 215 real singleton patients with an established diagnosis from the Deciphering Developmental Disorders (DDD) project *(35)*. The DDD dataset includes HPO phenotypes (a median of 7 per patient), exome data in variant call format (VCF), and the causative gene for each patient (1 per patient). AMELIE’s goal was to rank the established causative gene at or near the top of its ranked list of candidate genes for each patient. Filtering for candidate causative variants resulted in a median of 163 variants in 127 candidate genes per patient (Supplementary Methods and Figure 1c).

#### AMELIE outperforms previous methods at ranking candidate causative genes

The set of 215 patients obtained from the DDD study was used to evaluate AMELIE against Exomiser *(14)*, Phenolyzer *(15)*, Phen-Gen *(16)*, eXtasy *(17)*, and PubCaseFinder *(18)*. The output of all methods, consisting of a list of ranked genes, was subset to the (median) 127 candidate genes that AMELIE used for each patient based on the filtering criteria previously described (Figure 2a). This ensured the fair evaluation of all gene ranking methods against the same set of genes.

**Figure 2.**
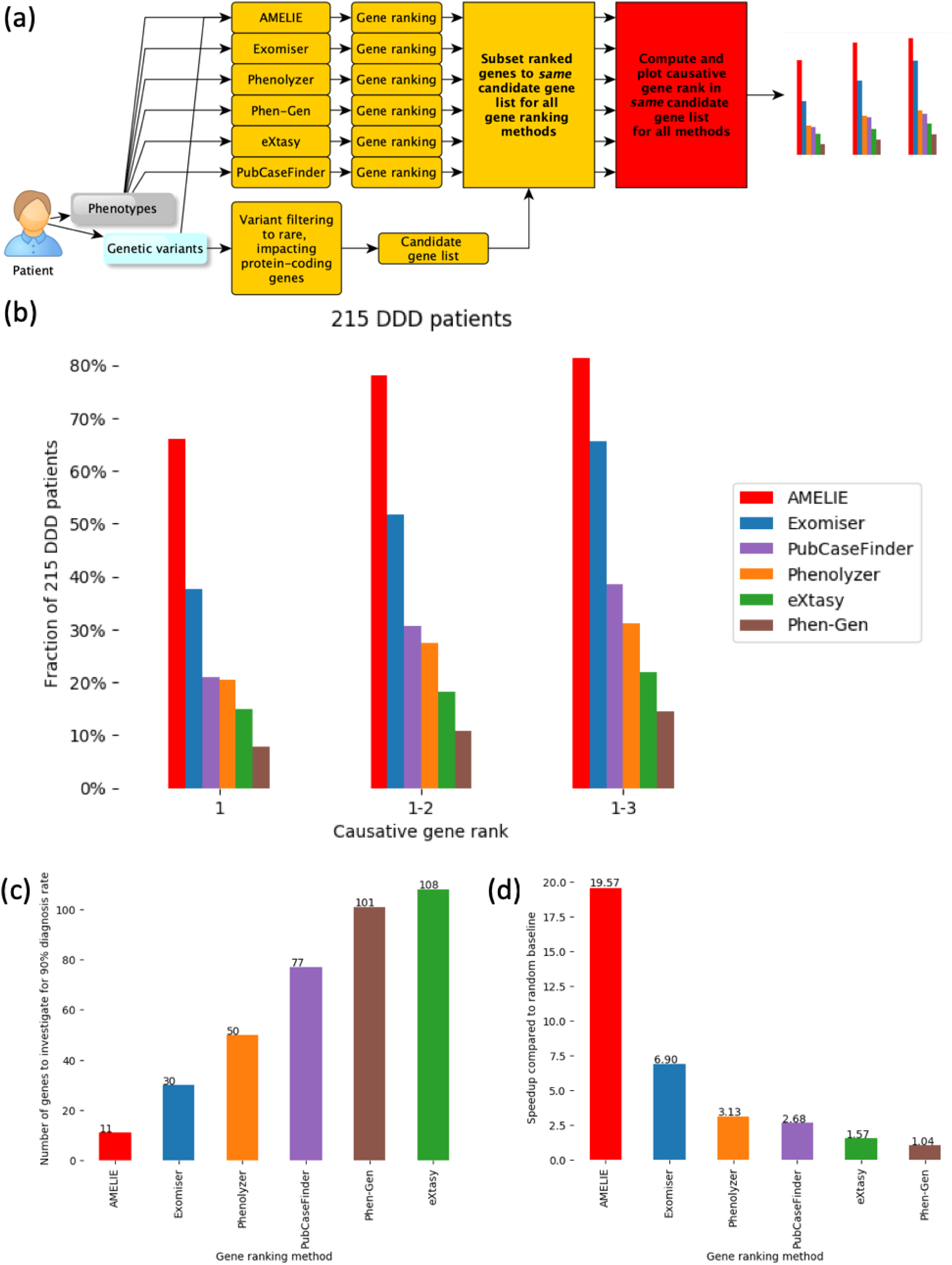
AMELIE patient causative gene ranking outperforms methods based on manually curated databases. **(a)** Evaluation scheme: The output gene ranking of all algorithms is subset to the same list of candidate genes AMELIE uses for its gene ranking to ensure a fair comparison. **(b)** Fraction of 215 DDD cases ranked as 1, 1-2 or 1-3 by six different tools. AMELIE ranks 66% of causative genes at number 1, nearly twice as many as the next best tool. **(c)** Rapid mode: By evaluating only up to AMELIE’s 11^th^ top ranked gene, one attains 90+% yield over DDD cases. The next best tool can guarantee this only by evaluating up to the 30^th^ gene. **(d)** In-depth mode: When evaluating each case, gene by gene, until a diagnosis is reached, AMELIE provides nearly 20x speed up over evaluation using random gene order. More than twice as fast as the next best tool. Figure S2 replicates this performance on two additional real patient cohorts.

AMELIE analyzed a median of 4,173 articles per patient. AMELIE ranked the causative gene at the very top in 142 (66%) out of 215 cases, and in the top 10 in 193 cases (89.7%). Other methods ranked the causative gene at the top between 38% of cases (Exomiser) and 8% of cases (Phen-Gen) (Figure 2b). AMELIE performed significantly better than all compared methods (all p-values ≤ 1.68*10^−9^; one-sided Wilcoxon signed rank test; see Supplementary Table S4 for all rankings).

#### Evaluating AMELIE’s top ranked 11 of 127 candidate genes already yields 90+% of diagnoses

Due to the large number of patients expected to be sequenced for Mendelian diagnosis *(36)*, one may want to set guidelines for rapid vs. in depth exome/genome analysis. Based on our 215 patients test set, AMELIE offers a diagnosis for 90% of diagnosable cases if one is willing to evaluate (in rapid mode) only up to the top 11 AMELIE ranked genes per case (9% of a median of 127 candidate causative genes). If using any of the other methods, the clinician would have to investigate between a median of 30 genes (Exomiser) and 108 genes per patient to arrive at the diagnosis in 90% of diagnosable cases (Figure 2c).

#### AMELIE needs 3-19x fewer patient-gene evaluations to diagnose all 215 cases

If the clinician uses AMELIE (in in-depth mode) to determine the order in which they evaluate their entire candidate gene list, one gene after the other, on the DDD set of 215 patients, they would evaluate a total of 735 gene-patient matches (summed across all patients) to arrive at the causative gene for all 215 patients. If the clinician went through the list of candidate genes in random order, they would evaluate an expected total of 14,383 gene-patient matches to arrive at the causative gene for all patients. By this metric, AMELIE improves diagnosis time by a factor of 19.6x over a random baseline. The next best tool, Exomiser, would require the clinician to read on 2,085 genes until arriving at the causative gene for all patients, an improvement of 6.9x over a random baseline. The performance of other methods ranges from a speedup of 3.13x to 1.04x (Figure 2d).

### Replication of AMELIE performance on 56 clinical cases from two sites

To test for result replication across data sources, we evaluated AMELIE using 56 singleton clinical cases seen by the Medical Genetics Service at Stanford Children’s Health and the Manton Center for Orphan Disease Research at Boston Children’s Hospital. Informed consent was obtained from all participants. Patient genotype and phenotype data were obtained from Stanford and the Manton Center Gene Discovery Core (Supplementary Methods).

#### AMELIE again outperforms all other methods at causative gene ranking

We performed a comparison of gene ranking performance using AMELIE against other methods as above for the DDD patients. AMELIE ranked the causative gene at the very top in 33 (59%) out of 56 cases, and in the top 10 in 50 cases (89%). Again, AMELIE significantly outperformed all other methods (all p-values ≤ 6.65*10^−3^; one-sided Wilcoxon signed rank test; see Supplementary Figure S2a and Supplementary Table S5 for all-method comparison).

#### AMELIE’s top ranked 9% candidate genes again contain 90% of diagnoses

AMELIE offered a diagnosis for 90% of patients in the test set of 56 Stanford and Manton patients if evaluating the top 15 candidate genes per patient (9% of a median of 172.5), thereby replicating its performance on the DDD set (Supplementary Figure S2b).

#### AMELIE needs 2-20x fewer patient-gene evaluations to diagnose clinical cases

Again, we calculated speedup of causative gene evaluation using AMELIE and other gene ranking methods compared to a baseline in which the clinician uses no automatic gene ranking methods. To arrive at the causative gene for each patient in the clinical test set from Stanford and Manton when using AMELIE, the clinician would evaluate 300 genes, compared to a baseline of 6,106 genes when evaluating genes in random order. Similar to the DDD patient test set, this results in a speedup of 20x, 2-20x fewer compared to other methods (Supplementary Figure S2c).

### Cross-validation of AMELIE causative gene ranking performance on simulated patients set

We tested AMELIE causative gene ranking performance on the 681 simulated patients. To do this, we split the set of simulated patients into 5 evenly sized chunks. In five round-robin iterations, we re-trained the AMELIE classifier using 4 of the 5 chunks of simulated patients, and evaluated the re-trained classifier’s causative gene ranking performance on the remaining fifth chunk of simulated patients (5-fold cross-validation). Combining the causative gene ranking results of all 5 cross-validation evaluations thus resulted in AMELIE causative gene ranks for 100% of the 681 simulated patients. AMELIE ranked the causative gene at the top in 621 cases (91%) and in the top 10 in 672 cases (99%), replicating previous results indicating far higher gene ranking performance on simulated patient data compared to real-world data *(12, 13, 15, 16, 37, 38)*. Compared to a random baseline, AMELIE speeds up causative gene discovery by 34.8x on simulated patients. The next best method, Exomiser, speeds up causative gene discovery by 17.9x and other methods perform worse (Supplementary Figure S3, Supplementary Table S6).

### AMELIE classifier utilizes its broad feature set to arrive at gene rankings

We ran multiple tests with modified AMELIE knowledgebases and AMELIE classifiers to dissect the relative contribution of different AMELIE components to its causative gene ranking performance.

#### Phenotypic match is the most important feature for most patients with high causative gene ranks

For all 175 real test patients with the causative gene ranked at the top, we investigated which features of the AMELIE contributed most to the high score of the causative gene. Overwhelmingly, for 149 (85%) of 175 real test patients, the feature contributing most to the high score was a high phenotypic match between the patient and the article. However, 14 out of a total 27 AMELIE classifier features (52%) occurred at least once within the 3 features contributing most to the top rank of a patient’s causative gene (Figure 3a and Supplementary Table S7).

**Figure 3.**
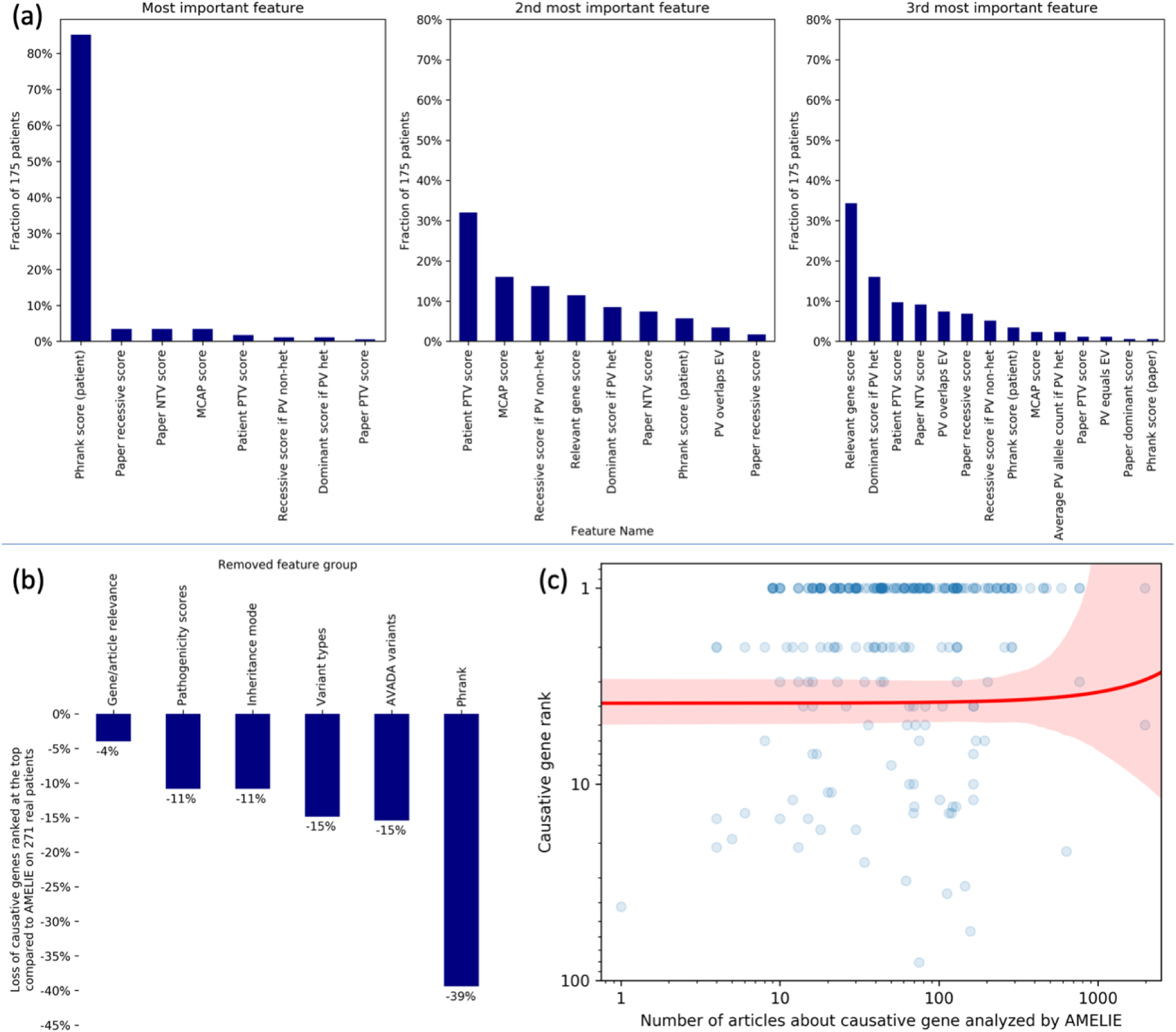
Investigating AMELIE’s gene ranking performance. **(a)** For each of the 175 patients with AMELIE causative gene rank 1, the 27 features to the AMELIE classifier were ranked by their contribution to the top-ranked article’s high score. The panels, left-to-right, show the fraction of patients for which certain features were ranked most, 2^nd^ most, or 3^rd^ most contributing. **(b)** Re-training the AMELIE classifier in round-robin fashion, each time omitting one of AMELIE’s 6 feature groups, finds all feature groups aid performance. **(c)** Each blue dot represents one of 271 real patients in this log-log plot. The red line is a linear regression line between number of articles about causative gene (x-axis) and causative gene rank (y-axis), with red denoting the 95% confidence interval. The slope of the regression line is not significantly different (Wald Test) from 0, suggesting AMELIE performs well independent of the number of papers it has read about a causative gene.

#### Removing features from the AMELIE classifier decreases performance

To measure how much AMELIE relied on certain feature groups, we re-trained the AMELIE classifier 6 times, each time dropping one of its 6 feature groups. With dropped-out features, the number of causative genes ranked at the top across the test set of 271 real patients shrank between 4% and 39% (Figure 3b and Supplementary Table S8).

#### Augmenting phenotype recognition in articles does not lead to increased performance

HPO cross-links some of its phenotype entries to other databases containing phenotype names. We utilized 19,949 cross-links available in HPO by augmenting HPO phenotype phrases with synonyms from UMLS *(39)*, MeSH *(40)*, and SNOMED-CT *(41)*, three databases containing phenotype names (Supplementary Methods). This augmentation of recognizable phenotype names increased the number of distinct recognized phenotypic abnormalities per article in the AMELIE knowledgebase from 9 to 22. However, on the main task of causative gene ranking, the augmented AMELIE knowledgebase did not perform better (169 instead of 175 patients had the causative gene ranked at the top).

#### AMELIE knowledgebase constructed from full text is up to 43% superior to title/abstract-only knowledgebase on clinical patients

To quantify the information gained from full text for the purpose of causative gene ranking compared to title/abstract-only data, we re-trained all AMELIE knowledgebase classifiers only on title/abstract data from PubMed. Subsequently, we constructed the AMELIE knowledgebase using title/abstract data from PubMed only and re-trained the AMELIE classifier using the title/abstract-only knowledgebase (Supplementary Methods). Title/abstract-based gene ranking performs significantly worse than ranking based on full text information across all 271 real test patients (p=1.87*10^−3^; one-sided Wilcoxon signed rank test). On the set of clinical patients from Stanford and Manton, 43% more patients had their causative genes ranked at the top using full-text based ranking compared to title/abstract-based ranking. Overall, title/abstract-based ranking put the causative genes of 133 of 271 real patients at the top, compared with 175 (+32%) of 271 real patients for full-text based AMELIE.

### AMELIE performance is not correlated with number of articles about causative gene

We investigated whether the number of articles about the causative gene in the AMELIE knowledgebase correlates with the causative gene rank by performing linear regression between the causative gene rank and number of articles analyzed for the causative gene. This revealed no significant correlation (p=0.85 that the slope of regression equals 0 according to a Wald Test with t-distribution of the test statistic; Figure 3c). On the 22 patients (8% of a total of 271 real test patients) with less than 10 papers analyzed for the causative gene, AMELIE had 10 causative genes ranked at the top (45%) compared to Exomiser, which ranked the causative gene at the top in 6 (27%) of these cases.

### AMELIE knowledgebase and AMELIE classifier work together to arrive at high causative gene ranks

We investigated the relative contribution of the AMELIE classifier and the AMELIE knowledgebase to AMELIE’s overall gene ranking performance.

#### AMELIE knowledgebase outperforms most up-to-date automatically curated knowledgebase

Multiple previous methods for text mining gene, disease, variant, and phenotype information from literature have been developed *(42–51)*, most of which curate limited information or information from comparatively small sets of articles. Of 3 additional efforts *(52–54)* attempting to curate gene-phenotype information from a broad set of articles in automatic or semi-automatic fashion, we focused on DisGeNET (the most recently updated such database), containing gene-phenotype relationships, disease-causing variants, and links to primary literature from PubMed. DisGeNET contains both automatically curated data and hand-curated data *(54)*.

We re-trained the AMELIE classifier using DisGeNET data (Supplementary Methods). Three different versions of DisGeNET information were used: **(1)** containing only data curated by DisGeNET’s BeFree automatic curation software, **(2)** containing only manually curated data in DisGeNET, **(3)** containing all suitable data in DisGeNET, both manually and automatically curated. All 3 versions of DisGeNET performed worse than AMELIE (all p-values ≤ 4.76*10^−23^, one-sided Wilcoxon signed rank test). Best-performing was the DisGeNET database with automatically curated data (68 causative genes ranked at the top, compared to 175 for original AMELIE; see Supplementary Table S9).

#### AMELIE classifier outperforms simple phenotype-based ranking

We replaced the AMELIE classifier (Figure 1c) with the Phrank *(11)* phenotypic match score to estimate the impact of the AMELIE classifier on overall AMELIE performance. For each of the 271 real test patients, we re-ranked articles about the candidate causative genes by computing the Phrank match score between patient phenotypes and phenotypes mentioned in the article. Candidate causative genes were then ranked by the Phrank score of the highest-ranked article about the gene. This resulted in ranking 94 (35%) of 271 real patients’ causative genes at the top, significantly worse compared to ranking based on the AMELIE classifier, which ranked 175 causative genes at the top (p=1.33*10^−11^, one-sided Wilcoxon signed rank test).

We conclude that the AMELIE knowledgebase and the AMELIE classifier work together to achieve AMELIE’s high causative gene ranking performance.

### Interactive and programmatic access to AMELIE-based literature analysis

AMELIE can be used through its web portal at https://AMELIE.stanford.edu to utilize AMELIE for patient analysis. The portal offers both an interactive interface (Supplementary Figure S4) and an application programming interface (API) that enables integrating AMELIE into any computer-assisted clinical workflow. The AMELIE knowledgebase will be updated every year. A pilot *(55)* of AMELIE has been running at this web address since August 2017, as a service to the community, using an AMELIE knowledgebase automatically curated from articles published until June 2016, and has since served many thousands of queries from 40+ countries.

## Discussion

We present AMELIE, a method for ranking candidate causative genes and supporting articles from the primary literature in patients with suspected Mendelian disorders. We show that AMELIE ranks the causative gene first (among a median of 127 genes) in 2/3 of patients, and within the top 11 genes in over 90% of 215 real patient cases. These results were closely replicated on a cohort of 56 clinical patients from Stanford Children’s Health and the Manton Center for Orphan Disease Research.

Mendelian disease diagnosis is a complex problem and clinicians or researchers can spend many hours evaluating a single case. With 5,000 diagnosable Mendelian diseases, caused by roughly 3,500 different genes, and manifesting in different subsets of over 13,000 documented phenotypes, manual patient diagnosis from the primary literature is highly labor intensive. Manually curated databases like OMIM, OrphaNet and HGMD take a step towards alleviating clinician burden by attempting to summarize the current literature. However, manual curation is growing ever more challenging as the literature about Mendelian diseases is increasing at an accelerating rate. Based on AMELIE analysis, the number of gene-phenotype relationships in Mendelian literature has been increasing by an average of 10.5% every two years since the year 2000 (Supplementary Figure S5). Because AMELIE is an automatic curation approach, requiring only an initial critical mass of human curated data to train on, it is not constrained by the bottleneck of on-going human curation. For example, of 117 top-ranked articles supporting the DDD patients where AMELIE ranked the test set causative gene at number 1, only 36 (31%) were cited in OMIM (as determined by a systematic Google search of omim.org; Supplementary Table S10). OMIM, a manually curated database, does not, of course, promise to capture all papers pertaining to any given disease gene, but an automated effort like AMELIE can.

Compared to existing gene ranking approaches, AMELIE replaces the notion of a fixed disease description (e.g., as a single set of phenotypes) with the notion of an article and the phenotypes described in it. This approach has multiple advantages. First, it is often fastest to convince a clinician about a diagnosis given an article directly describing the disease, which often includes disease information such as patient images and related literature. Additionally, with considerable phenotypic variability in Mendelian diseases *(56)*, matching patients to specific reports in the literature is conceptually more helpful for definitive diagnosis than matching to a disease, which is effectively a compendium of previously described patient phenotypes. Biomedical literature is not guaranteed to contain the full set of phenotypes known to be associated with a disease, and AMELIE makes no claim about capturing this full set. Rather, AMELIE focuses on causal gene ranking, using its knowledgebase, and as we show, it already does it to great practical utility. Certain articles about Mendelian diseases may mention a very small number of phenotypes (or none at all) and just mention disease and causative genes. While this situation does not appear to be very common in practice (as seen by the good performance of AMELIE), the problem could be alleviated by automatically parsing disease names from such articles and associating diseases with manually curated phenotype information using resources such as HPO. Natural language processing approaches can also be used to read additional texts as well, even sources like electronic medical notes *(19, 20)*.

Understanding the impact of hundreds of thousands of variants in thousands of different genes against a body of knowledge of millions of peer reviewed papers that is ever expanding is a challenging task. Because a diagnosis shapes the future management of a patient, there must always be a human expert approving every diagnosis. But the sheer number of patients that can benefit from a molecular diagnosis and our intention to sequence millions of them in the next few years absolutely necessitate automating as much as possible of the diagnostic process, to potentiate rapid, affordable, reproducible and accessible clinical genome-wide diagnosis. As such, along with complementary medical record parsing tools *(19, 20)*, AMELIE provides an important step on the road to integrating personal genomics into standard clinical practice.

## Acknowledgments

We thank Dr. Kai Wang for his assistance with ANNOVAR and Erich Weiler for support and guidance. We thank Paul McDonagh and Thomas Defay, as well as the members of the Bejerano lab, for technical advice and helpful discussions. We thank E. Kravets, J. Buckingham and K. MacMillen for assistance with obtaining patient data. We thank all data sources used in AMELIE, including HPO, Ensembl, HGNC, UniProt, OMIM, ClinVar, HGMD, PubTator, ExAC, pLI, RVIS, M-CAP, and the GWAS catalog. We thank the European Genome-Phenome Archive ***(57)*** (EGA) and the Deciphering Developmental Disorders ***(35)*** (DDD) project for sharing some data. The DDD study presents independent research commissioned by the Health Innovation Challenge Fund [grant number HICF-1009-003], a parallel funding partnership between the Wellcome Trust and the Department of Health, and the Wellcome Trust Sanger Institute [grant number WT098051]. The views expressed in this publication are those of the author(s) and not necessarily those of the Wellcome Trust or the Department of Health. The study has UK Research Ethics Committee approval (10/H0305/83, granted by the Cambridge South REC, and GEN/284/12 granted by the Republic of Ireland REC). Deidentified DDD data was obtained through EGA. The research team acknowledges the support of the National Institute for Health Research, through the Comprehensive Clinical Research Network.

## Funding

All computational work was funded in part by a Bio-X SIGF fellowship to J.B., DARPA (C.R., G.B.), the Stanford Pediatrics Department (J.A.B., G.B.), a Packard Foundation Fellowship (G.B.), a Microsoft Faculty Fellowship (G.B.), NHGRI U41HG002371-15 (M.H.) and the Stanford Data Science Initiative (G.B., C.R.). Manton Center sequence analysis and diagnosis was supported by NIH 1U54HD090255 IDDRC Molecular Genetics Core grant. The content is solely the responsibility of the authors and does not necessarily represent the official views of the National Institutes of Health.

## Author contributions

J.B. and G.B. designed the study and analyzed results. J.B. implemented the text mining software, website and associated databases. M.H. authored PubMunch. J.B., E.H.S., H.G., A.M.W., M.E.D., K.A.J., and C.A.D. wrote and improved software tools that were used for genotype and phenotype analysis. C.A.D. analyzed EGA data. A.J.R. wrote parts of the gene and phenotype identification. P.D.S. and D.N.C. curated the HGMD data. C.R. provided text mining guidance. The Manton Center and A.H.B. provided Manton patient data from the Gene Discovery Core. J.A.B. provided guidance on clinical aspects of study design, testing set construction and interpretation of results. J.B. and G.B. wrote the manuscript. G.B. supervised the project. All authors commented on and approved the manuscript.

